# Culture influences audiovisual emotion perception in early sensory areas

**DOI:** 10.1101/2020.10.10.332437

**Authors:** Akihiro Tanaka, Sachiko Takagi, Tokiko Harada, Elisabeth Huis In ’t Veld, Beatrice de Gelder, Yuki Hamano, Ken-ichi Tabei, Norihiro Sadato

## Abstract

Emotion perception from facial and vocal expressions is a multisensory process critical for human social interaction. When asked to judge emotions by attending to either face or voice, the accuracy was higher when facial expressions are congruent with vocal expressions than when they are incongruent. This congruency effect was shown to be affected by cultural background. Here we conducted functional MRI alongside a multisensory emotion perception task involving Japanese and Dutch participants. They were presented with movies in which congruent or incongruent emotions were expressed through faces and voices. The participants were asked to judge the emotion of either the face or the voice. Consistent with previous studies, behavioral results showed an interaction between group and task. fMRI results revealed that during voice-based judgement, congruency effects of the primary visual cortex, by means of the task related activity of congruent stimuli subtracted by that of incongruent stimuli, were more prominent in the Dutch group than in Japanese group. Finally, behavioral and neural congruency effects of the primary visual cortex were positively correlated only in the Dutch group. Taken together, our results show that culture affects the activities of early sensory areas in multisensory perception from facial and vocal expressions.

## Introduction

Accurate perception of other people’s emotions is a prerequisite for successful social interactions. Emotions are expressed through different sensory channels, such as the face, body, and voice, and are judged by integrating information from these channels in natural settings. The literature has shown interactions in emotion perceptions from facial and vocal expressions (Collignon et al., 2008; de Gelder & Bertelson, 2003; de Gelder et al., 1999; de Gelder & Vroomen, 2000; Koizumi et al., 2011; Liu et al., 2015; Massaro & Egan, 1996; Takagi et al., 2015; Tanaka et al., 2010). For instance, de Gelder and Vroomen (2000) asked participants to judge the emotion of congruent and incongruent face-voice stimuli expressing two opposing emotions (happiness and sadness) by instructing participants to attend to a specific modality. The results showed that accuracy of emotion perception was higher for congruent stimuli than for incongruent stimuli (i.e., congruency effect).

This congruency effect, which is a measure of the cross-modal bias, was shown to be affected by cultural background (Tanaka et al. 2010). Tanaka et al. (2010) investigated cultural differences between Japanese and Dutch people in terms of the multisensory perception of emotion. A face and a voice, expressing either congruent or incongruent emotions, were presented in each trial (e.g., a happy face was presented with an angry voice, in the incongruent case) using the immediate cross-modal bias paradigm (Bertelson & de Gelder, 2004). Participants were instructed to judge the emotion expressed in one of the two sources (face or voice) and to ignore the other source. Congruency effects were compared between Japanese and Dutch participants. The results demonstrated that when the face and the voice did not represent the same emotion, Japanese participants weighted cues in the voices more than the Dutch participants. These findings provided first evidence that culture affects the multisensory integration of affective information.

Using a similar paradigm, Liu et al. (2015) compared the multisensory emotion perceptions between North American and Chinese people as indexed by both behavioral and electrophysiological measures. Results indicated that while both groups were sensitive to emotional differences between channels (with lower accuracy and higher N400 amplitudes for incongruent face-voice pairs), there were marked between-group differences in how incongruent facial or vocal cues affected accuracy and N400 amplitudes, with North American participants showing greater interference from irrelevant faces than Chinese participants. These results show temporal properties of neural processing of multisensory emotion perception and suggest that culture modulates semantic processing of the stimuli as indexed by N400.

Neuroimaging studies have investigated the neural basis of audiovisual integration of emotional information (de Gelder et al., 1999; Dolan et al., 2001; Ethofer et al., 2006, 2013; Jansma et al., 2014; Klasen et al., 2011; Kreifelts et al., 2007, 2009, 2010; Muller et al, 2011; Pourtois et al., 2005). Specifically, several studies have reported brain activities when contrasting audiovisual (face and voice) to visual (face-only) and/or auditory (voice-only) conditions. These studies reported stronger activation in the left middle temporal gyrus (Pourtois et al., 2005), fusiform gyrus (Kreifelts et al., 2010), amygdala (Kreifelts et al., 2010), thalamus (Kreifelts et al., 2007, 2010), and posterior superior temporal sulcus (pSTS; Pourtois et al.,2005; Ethofer et al., 2013; Kreifelts et al., 2007, 2009, 2010). Other studies used facial and vocal stimuli expressing congruent and incongruent emotions (e.g., Dolan et al., 2001; Ethofer et al., 2006; Klasen et al., 2011; Muller et al., 2011). Dolan et al. (2001) showed stronger activations in amygdala and fusiform area to fearful faces when the face expresses congruent emotion with a simultaneously presented voice. Muller et al. (2011) reported that the incongruence of emotional valence between faces and sounds led to increased activation in the middle cingulate cortex, right superior frontal cortex, right supplementary motor area as well as the right temporoparietal junction.

How are the perceived emotions represented in the brain? Emotional representation is considered to be supra-modal. Literature has shown that the heteromodal limbic and paralimbic regions including ventral posterior cingulate and amygdala represent a supramodal representation of emotions (e.g., Klasen et al., 2011; Scott et al., 1997), as well as cortical regions including medial prefrontal cortex, left superior temporal sulcus (Peelen et al., 2010), and orbitofrontal cortex (Chikazoe et al., 2014). Previous studies have reported cultural effects on cognitive processing of negative facial expressions in the amygdala, a part of the limbic system that is the key structure for emotional processing (Chiao et al., 2008; Harada et al., 2020). Recent studies provide converging evidence that early sensory cortices also code the valence of socio-emotional signals (Miskovic & Anderson, 2018; Shinkareva et al., 2014). Miskovic and Anderson (2018) argue that these modality-general and modality-specific systems may play differential roles in emotion judgement. Thus, in order to identify the locus of crossmodal interaction in emotion perception, it is necessary to examine neural activities in early sensory cortices and to examine cultural effects.

In this study, we investigated the neural basis of cultural differences in emotion perception from multisensory signals. Specifically, we focused on the roles of early sensory areas in the multisensory perception of emotion. On the basis of literature using the cross-modal bias paradigm, we predicted that behavioral performance would be worse in incongruent conditions than in congruent conditions in both face and voice tasks. On the basis of the findings of cultural differences (Tanaka et al., 2010), we predicted that this congruency effect would be different depending on the culture of participants. On the basis of the literature showing early influences of cross-modal signals using simple audiovisual stimuli (Watkins et al., 2006, 2007), we predicted that congruency between facial and vocal emotions would affect the activity of the primary sensory areas (i.e., primary auditory and visual cortices). On the basis of the literature on the valence coding in early sensory cortices, we also predicted that there are cultural differences in the activities of early sensory cortices.

## Methods

### Participants

Fifty-three participants (28 Japanese and 25 Dutch) were recruited as paid volunteers for the fMRI experiments. Eighteen participants (8 Japanese and 10 Dutch) were excluded due to large amount of head movements and high rates of response errors during the fMRI session, leaving 35 participants for the final analysis: 20 Japanese (11 females, 9 males; mean age = 24.35 ± 4.77 (*SD*) years) living in Japan and 15 Dutch (9 females, 6 males; mean age = 25.53 ± 3.96 (*SD*) years) living in the Netherlands. All participants had normal or corrected-to-normal vison. The participants were all right-handed and had no neurological disorders. Written informed consent was obtained from each participant before the experiment. This study was approved by the Ethical Committee of the National Institute for Physiological Sciences, Tokyo Woman’s Christian University, Japan, and Tilburg University, Netherlands.

### Experimental stimuli

In the emotion judgment task, we used audiovisual stimuli that were created from videos of two Japanese and two Dutch female speakers uttering four short fragments with neutral linguistic meaning (e.g., “What is this?”, “Good bye” and so on). The speakers uttered the short fragments in their native language with happy or angry emotions. Happy or angry facial expressions were combined with happy or angry vocal expressions, resulting in a total of 64 bimodal stimuli (two speakers × four fragments × two facial expressions × two vocal expressions × two languages). Consequently, the bimodal stimuli set consisted of 32 emotionally congruent (e.g., a happy face with a happy voice) and 32 emotionally incongruent stimuli (e.g., a happy face with an angry voice). We also used 64 unimodal stimuli: 32 face stimuli (i.e. happy or angry faces) and 32 voice stimuli (i.e. happy or angry voices). A 95% random dynamic noise was added to the visual stimuli to avoid a ceiling effect and to match participants’ performances between face-only and voice-only trials during an emotional judgment task with unimodal stimuli (Tanaka et al., 2010).

### Experimental procedure

To avoid any cross-site scanner effect, all MRI and behavioral data (i.e. both of Japanese and Dutch data) were acquired using the same MRI machine (Allegra; Siemens, Erlangen, Germany) located in the National Institute for Physiological Sciences in Japan. Before the fMRI session, participants were given detailed instructions regarding the task procedure. To familiarize the participants with the task, they were presented with examples of stimuli in a practice session. No stimulus in the practice session was used in the later fMRI session. All stimuli were presented using the Presentation 14.8 software (Neurobehavioral Systems, Albany, CA, USA) running on a personal computer (Dimension 9200; Dell Computer, Round Rock, TX, USA). Using a liquid crystal display (LCD) projector (DLA-M200L; Victor, Yokohama, Japan), the visual stimuli were projected onto a half transparent viewing screen located behind the head coil of the magnetic resonance imaging (MRI) scanner. Participants viewed the stimuli via a tilted mirror attached to the head coil. The spatial resolution of the projector was 1024 × 768 pixels with a 60-Hz refresh rate. The distance between the screen and the eyes was approximately 60 cm, and the visual angle was 18.9° × 14.2° (vertical). Auditory stimuli were presented via MR-compatible headphones (Hitachi, Yokohama, Japan).

The fMRI session consisted of two task conditions: the face and voice task conditions. The participants were asked to complete two task runs for each task condition, resulting in a total of four task runs. In the face task runs, audiovisual stimuli (visual angle 18.9° × 14.2°) or face stimuli (visual angle 18.9° × 14.2°) were presented on the screen for 2.0 seconds, followed by a white fixation cross (visual angle 0.6° × 0.6°) on a black background for 0.5 seconds. When response alternatives (i.e. happy and angry) were presented for 2.0 seconds, the participants were required to categorize the emotion on the face as happy or angry while ignoring the voices. In the voice task runs, audiovisual stimuli or voice stimuli were presented, followed by a white fixation cross (visual angle 0.6° × 0.6°) on a black background for 0.5 seconds. When response alternatives were presented for 2.0 seconds, the participants were required to categorize the emotion on the voice as happy or angry while ignoring the face. They responded by pressing a button with their right index or middle finger during judgment periods through the task runs. The participants were instructed to set accuracy above speed.

We used an event-related design to minimize habituation and learning effects. A face task run included 48 task trials (i.e. 32 audiovisual and 16 face-only trials) and 16 null trials (i.e. a baseline condition) in which a white fixation instead of a video was presented for 4.0 seconds. A voice task run included 48 task trials (i.e. 32 audiovisual and 16 voice-only trials) and 16 null trials (i.e. a baseline condition). The stimuli were presented in a pre-defined randomized order in each task run. The order of the task conditions (i.e. face and voice task conditions) and the two task runs of each task condition were counterbalanced across participants.

### MRI data acquisition

All images were acquired using a 3-Tesla MR scanner (Allegra; Siemens, Erlangen, Germany). An ascending T2*-weighted gradient-echo echo-planer imaging (EPI) procedure was used for functional imaging to produce 42 continuous transaxial slices covering the entire cerebrum and cerebellum (time echo [TE], 30 ms; flip angle, 80°; field of view [FoV], 192 mm; 64 × 64 matrix; voxel dimensions, 3.0 × 3.0 mm in plan, 3.5 mm slice thickness with 15% gap). A “sparse sampling” technique was used to minimize the effects of image acquisition noise on task performance. Repetition time (TR) between two successive acquisitions of the same slice was 4500 ms. Cluster volume acquisition time was 2000 ms, leaving a 2500-ms silent period, during which an audiovisual, face, or voice stimulus was presented. Oblique scanning was used to exclude the eyeballs from the images. Each run consisted of a continuous series of 115 vol acquisitions, resulting a total duration of 8 min 38 sec. A high-resolution anatomical T1-weighted image (MPRAGE; TR = 2.5 s, TE = 4.38 ms, flip angle = 8 degrees, 256 × 256 matrix, 192 slices, voxel size = 0.75 mm × 0.75 mm × 1.0 mm) was also acquired for each participant. The total duration of the experiment was around 90 min for each participant.

### Data analysis

#### Behavioral performance

To examine whether audiovisual consistency of stimuli affects accuracy in both face and voice tasks regardless of culture, an analysis of variance (ANOVA) of the Task (face or voice) × Audiovisual consistency of stimuli (congruent or incongruent) within subjects was conducted. Then, to examine the general cross-modal bias (Bertelson & de Gelder, 2004), a mixed-factor ANOVA of Group (Japanese or Dutch) × Task (face or voice) was conducted on congruency effects, which were calculated by subtracting the mean accuracy in the incongruent condition from the mean accuracy in the congruent condition. The analysis was carried out using SPSS version 24.0 software (IBM Armonk, NY, USA).

#### fMRI data analysis

The MRI data were analyzed using SPM8 software (Wellcome Department of Imaging Neuroscience, London, UK) implemented in MATLAB (MathWorks, Sherborn, MA, USA). The first two EPI images of each task run were discarded because of unsteady magnetization, and the remaining 113 EPI images per run (a total of 452 EPI images per participant) were subjected to the following data analysis. All EPI images were spatially realigned to the mean EPI image to correct for head motion. The T1-weighted anatomical image was segmented into gray and white matters as well as CSF and reconstructed with a signal inhomogeneity correction. After the T1 weighted anatomical image was coregistered onto the mean EPI image, all EPI images were normalized to the MNI space (Montreal Neurological Institute [MNI] template) using a transformation matrix obtained from the normalization process of the T1 weighted anatomical image of each individual participant to the MNI template. The normalized and resliced images were then spatially smoothed with a Gaussian kernel of 8 mm (full width at half-maximum; FWHM) in the X, Y, and Z axes.

After preprocessing, individual analysis of the EPI images obtained for each participant was conducted using a general linear model. During the first level analysis, the following conditions were separately modeled as regressors: 1) congruent conditions of Japanese stimuli (JCon), 2) incongruent conditions of Japanese stimuli (JIncon), 3) congruent conditions of Dutch stimuli (DCon), 4) incongruent conditions of Dutch stimuli (DIncon), 5) face-only conditions of Japanese stimuli (JFace), 6) face-only conditions of Dutch stimuli (DFace), 7) voice-only conditions of Japanese stimuli (JVoice), 8) voice-only conditions of Dutch stimuli (DVoice) and 9) null conditions. Each trial was modeled with a 4.5 second duration (i.e., from the onset of a stimulus to the end of a judgment period) convolving a hemodynamic response function. In addition, six regressors for movement parameters obtained in the realignment process were entered in the design matrix. An additional regressor of the mean signal from the cerebrospinal fluid (CSF) was also included in the design matrix. High-pass filters (128 s) were applied to the time-series data. An autoregressive model was used to estimate the temporal autocorrelation. The signals of images were scaled to a grand mean of 100 overall voxels and volumes within each run.

At the second-level analysis, in order to examine how culture might affect neural basis of multisensory perception of emotion, we primarily focused on the effect of culture on emotion perception in congruent and incongruent conditions by using a 2 × 2 factorial design: face-voice combinations (congruent or incongruent) and participant cultural groups (Japanese or Dutch). The four contrast images from each participant were subjected to the subsequent group analysis: [(JCon + DCon) - null] in the face task condition, [(JIncon + DIncon) - null] in the face task condition, [(JCon + DCon) - null] in the voice task condition, and [(JIncon + DIncon) - null] in the voice task condition. To examine the neural basis of cultural differences in emotion perception and the roles of early sensory areas in multisensory perception of emotion, we conducted a ROI analysis by extracting BOLD signals from four pre-defined ROIs (i.e. bilateral BA 17 and BA 41) based on WFU_PickAtlas, using MarsBaR version .44 (http://marsbar.Sourceforge.net).

## Results

### Behavioral performance

To examine whether audiovisual consistency of stimuli affects accuracy in both face and voice tasks regardless of culture, an ANOVA of Task (face or voice) × Audiovisual consistency of stimuli (congruent or incongruent) within subjects was conducted (Fig 1). The main effect of task, F (1, 34) = 20.56, p < .001, η_p_^2^ = .37, the main effect of audiovisual consistency of stimuli, F (1, 34) = 91.12, p < .001, η_p_^2^ = .73, and the two-way interaction between task and audiovisual consistency, F (1, 34) = 4.70, p < .05, η_p_^2^ = .12, were significant. Simple main-effects analyses showed that accuracy was higher for the congruent condition than for the incongruent condition in both face tasks, F (1, 34) = 41.85. p < .001, η_p_^2^ = .55, and voice tasks, F (1, 34) = 91.86, p < .001, η_p_^2^ = .73. Thus, the results show that the audiovisual consistency of stimuli affects accuracy.

**Fig 1.**
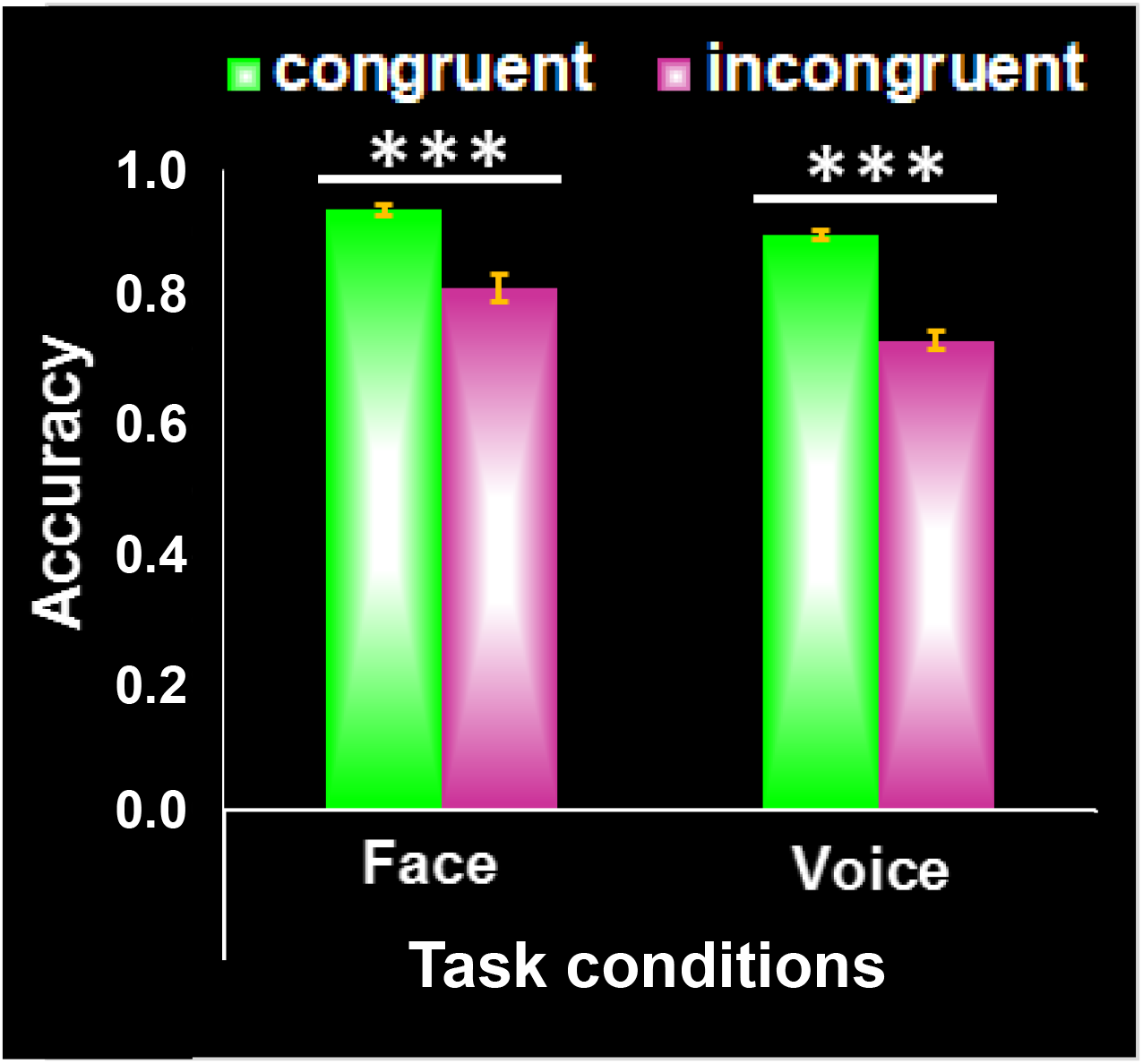
Mean accuracy of the congruent and incongruent conditions in the face and voice tasks. Error bars represent standard errors. Asterisks indicate significant differences between conditions (***p < .001).

Next, to examine the cross-modal bias in behavioral data, a mixed-factor ANOVA of Group (Japanese or Dutch) × Task (face or voice) was conducted on congruency effects (Fig 2). The main effect of task, F (1, 33) = 6.51, p < .05, η_p_^2^ = .17, and the two-way interaction between task and group, F (1, 33) = 4.41, p < .05, η_p_^2^ = .12, were significant. Simple main-effects analyses showed that the congruency effect was not different between face tasks and voice tasks in Japanese participants, F (1, 33) = 0.12, n.s., η_p_^2^ = .00, but was larger for voice tasks than for face tasks in Dutch participants, F (1, 33) = 9.46, p < .005, η_p_^2^ = .50.

**Fig 2.**
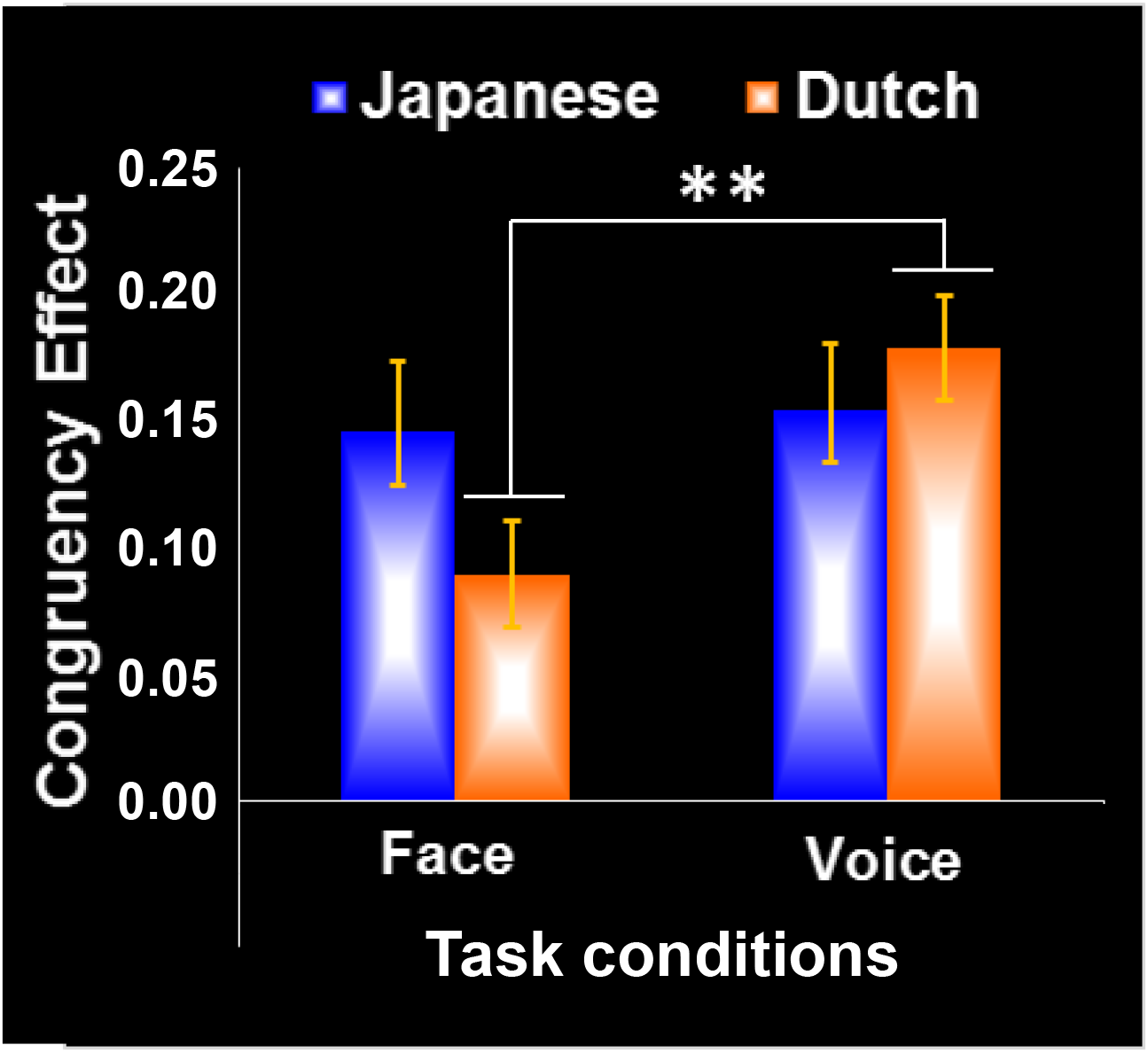
Congruency effects (i.e. mean accuracy in the congruent condition minus mean accuracy in the incongruent condition) in the face and voice tasks for the Japanese and Dutch participant groups. Error bars represent standard errors. Asterisks indicate significant differences between tasks in Dutch group (**p < .01).

### fMRI results

To examine whether audiovisual consistency affects the activity of the primary sensory areas and whether there might be cultural differences, we report the activities in primary sensory areas to emotionally congruent and incongruent audiovisual stimuli, and compare the congruency effects in the primary sensory cortices (i.e., task related activity of congruent stimuli subtracted by that of incongruent stimuli) between cultures.

To test whether audiovisual consistency might affect the activity of the primary sensory cortex (BA17L/R and BA41L/R), we conducted an ROI analysis in which an ANOVA of Task (face or voice) × Audiovisual consistency of stimuli (congruent or incongruent) within subjects was conducted on the effect size extracted from each of the four hypothesized regions (Fig 3). In BA17L, the main effect of audiovisual consistency, F (1, 34) = 4.89, p < .05, η_p_^2^ = .13, and the two-way interaction between task and audiovisual consistency, F (1, 34) = 5.41, p < .05, η_p_^2^ = .14, were significant. A simple main-effects analysis showed that the brain activity in BA17L was not different between the congruent condition and the incongruent condition in voice tasks, F (1, 34) = .38, n.s., η_p_^2^ = .01, but were larger for the congruent condition than for the incongruent condition in face tasks, F (1, 34) = 10.59, p < .005, η_p_^2^ = .24. In BA17R, the main effect of task, F (1, 34) = 6.07, p < .05, η_p_^2^ = .15, and the main effect of audiovisual consistency, F (1, 34) = 10.20, p < .005, η_p_^2^ = .23 were significant. In addition, the two-way interaction between task and audiovisual consistency, F (1, 34) = 3.77, p < .10, η_p_^2^ = .10, was marginally significant. A simple main-effects analysis showed that the brain activity in BA17R was slightly different between the congruent condition and the incongruent condition in voice tasks, F (1, 34) = 3.81, p < .10, η_p_^2^ = .10, but were larger for the congruent condition than for the incongruent condition in face tasks, F (1, 34) = 14.11, p < .005, η_p_^2^ = .29. The results suggest that audiovisual consistency affected brain activity in the bilateral primary visual cortices, especially in the face task condition.

**Fig 3.**
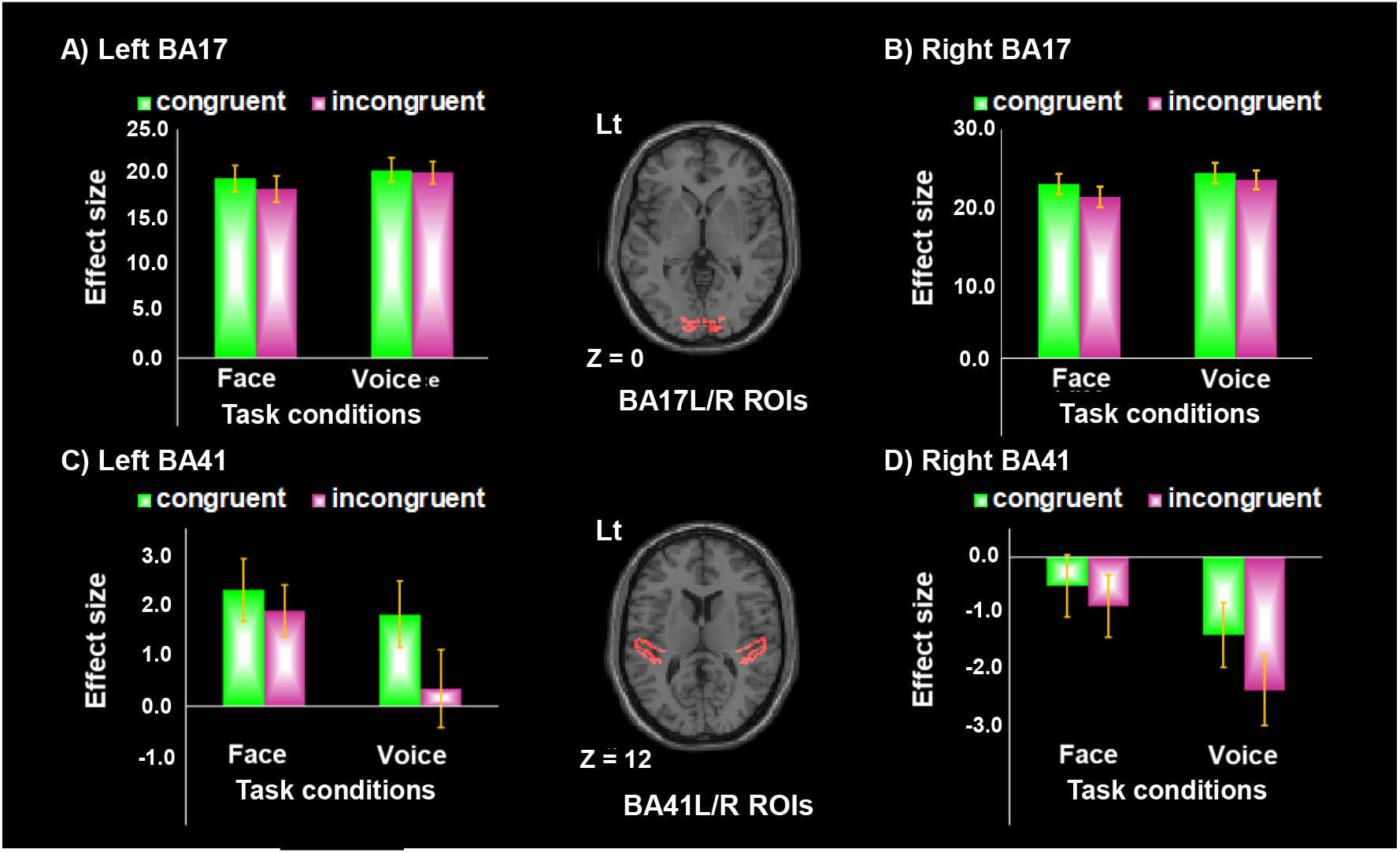
Activations in the congruent (green bars) and incongruent conditions (red bars) for the face and voice tasks within the bilateral primary visual (i.e. BA17) and auditory cortices (i.e. BA41). Effect sizes were extracted from Marsbar ROIs (shown in the middle column). The activities in the left and right BA17s were shown in (A) and (B), respectively. In both (A) and (B), activities were larger when facial and vocal emotions were congruent than when they were not congruent in the face task. The activities in the left and right BA41s were shown in (C) and (D), respectively. In both (C) and (D), activities were larger when facial and vocal emotions were congruent than when they were not congruent in the voice task. BA17L, the left BA17; BA17R, the right BA17; BA41L, the left BA41; BA41R, the right BA41.

In BA41L, the main effect of audiovisual consistency, F (1, 34) = 13.69, p < .001, η_p_^2^ = .29, and the two-way interaction of task and audiovisual consistency, F (1, 34) = 7.34, p < .05, η_p_^2^ = .18, were significant. A simple main-effects analysis showed that the brain activity in BA41L was not different between the congruent condition and the incongruent condition in face tasks, F (1, 34) = 1.87, n.s., η_p_^2^ = .05, but were larger for the congruent condition than for the incongruent condition in voice tasks, F (1, 34) = 18.82, p < .001, η_p_^2^ = 36. In BA41R, the main effect of task, F (1, 34) = 7.98, p < .01, η_p_^2^ = .19, and the main effect of audiovisual consistency, F (1, 34) = 13.56, p < .005, η_p_^2^ = .29, were significant. The two-way interaction between task and audiovisual consistency, F (1, 34) = 3.88, p < .10, η_p_^2^ = .10, was marginally significant. A simple main-effects analysis showed that the brain activity in BA41R was not different between the congruent condition and the incongruent condition in face tasks, F (1, 34) = 2.07, n.s., η_p_^2^ = .06, but were larger for the congruent condition than for the incongruent condition in voice tasks, F (1, 34) = 17.18, p < .001, η_p_^2^ = .34. These results suggest that audiovisual consistency affected brain activity in the bilateral primary auditory cortices, especially in the voice task condition.

To examine whether there might be cultural differences in congruency effects on brain activities in the primary sensory cortices (BA17 L/R and BA41 L/ R), a mixed-factor ANOVA of Group (Japanese or Dutch) × Task (face or voice) was conducted on congruency effects calculated from the effect size (i.e. a subtraction of effect size in the incongruent condition from effect size in the congruent condition) extracted from each of the four hypothesized regions (Fig 4). In BA17L, the main effect of task, F (1, 33) = 6.55, p < .05, η_p_^2^ = .17, was significant. In BA17R, the main effect of group, F (1, 33) = 5.33, p < .05, η_p_^2^ = .14, and the main effect of task, F (1, 33) = 4.67, p < .05, η_p_^2^ = .12, were significant. In both BA17L, F (1, 33) = 2.27, n.s., η_p_^2^ = .06, and BA17R, F (1, 33) = 2.12, n.s., η_p_^2^ = .06, a two-way interaction of task and group was not significant. To further examine a potential cultural effect on congruency effects in each task condition in detail, we conducted a simple main-effects analysis. The results showed that the congruency effects in brain activity were larger in Dutch than in Japanese in both BA17L, F (1, 33) = 8.57, p < .01, η_p_^2^ = .21, and BA17R, F (1, 33) = 8.26, p < .01, η_p_^2^ = .20, especially in the face task condition, but not in the voice task condition.

**Fig 4.**
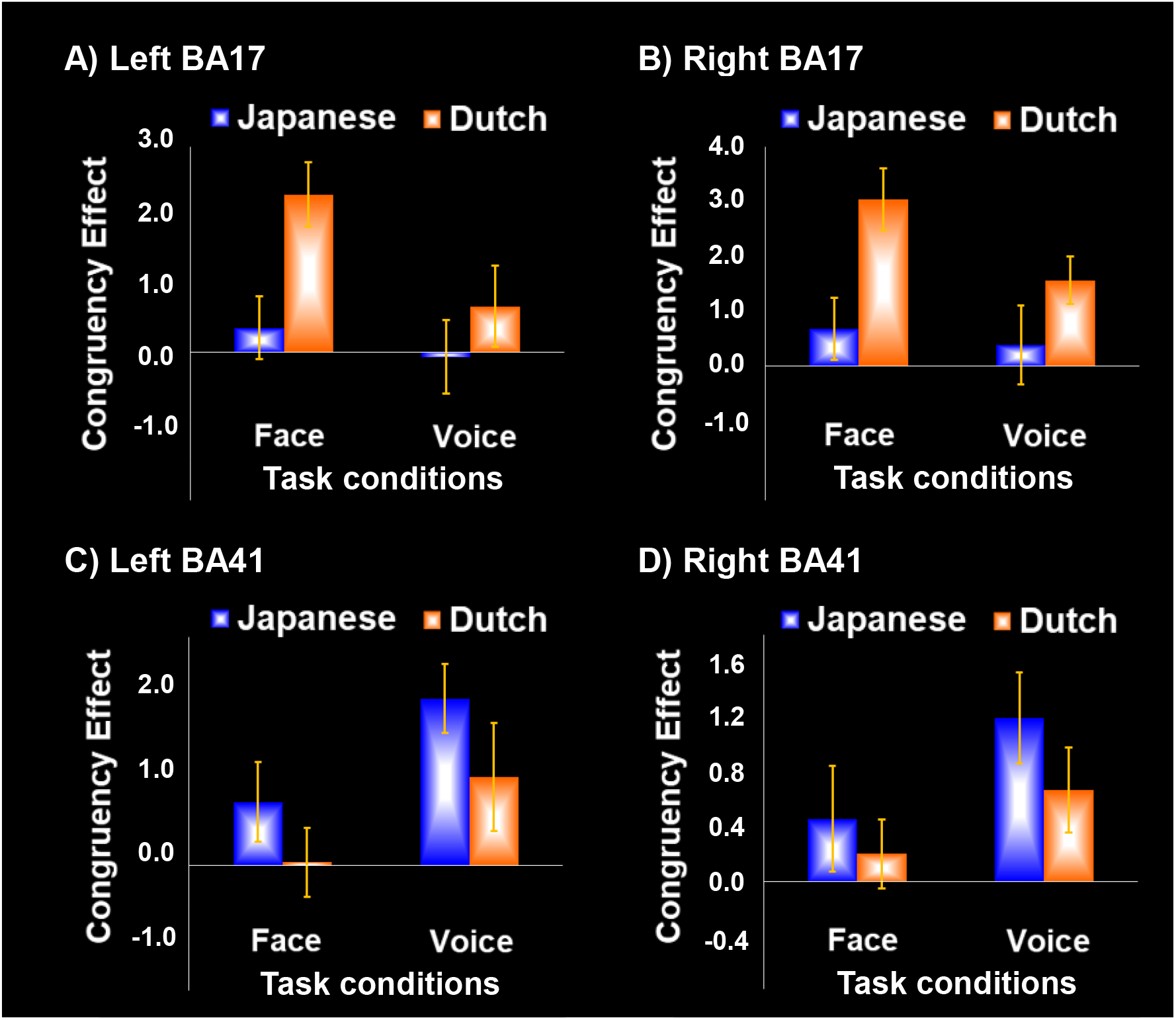
Congruency effects on brain activities within the primary visual (BA17) and auditory cortices (BA41) in the Japanese and Dutch groups. Effect sizes were extracted from Marsbar ROIs shown in the Figure 3, and the congruency effect on brain activities was calculated by subtracting the effect size of the incongruent condition from the that of the congruent condition. The congruency effects in the left and right BA17s are shown in (A) and (B), respectively. The congruency effects in the left and right BA41s were shown in (C) and (D), respectively. In the bilateral BA17s, the congruency effects on the brain activities are larger in the Dutch than Japanese group in the face task.

In BA41L, only the main effect of task, F (1, 33) = 6.80, p < .05, η_p_^2^ = .17, was significant. In BA41R, the main effect of task, the main effect of group, and the two-way interaction between task and group were not significant. In addition, as described above, a simple main-effects analysis was conducted. Results showed that the congruency effect in brain activity in both face and voice tasks were not different between Dutch and Japanese in BA41L/R.

### Pearson’s correlation analysis of behavioral data and imaging data in face task

As cultural differences were observed in brain activities for the face task condition, Pearson’s correlation analyses on congruency effects of behavioral and fMRI data during the face task were conducted in the primary sensory cortices (BA17 L/R and BA41 L/ R) (Fig 5). There was a positive correlation between the two variables in Dutch in BA17L, r = 0.62, p < .05, and BA17R, r = 0.54, r < .05. No significant correlation was found in the primary auditory cortex in Dutch participants. In contrast, there was a marginally significant negative correlation between the two variables in Japanese in BA41L, r = −.39, p < .10. No significant correlation was found in the primary visual cortex in Japanese participants. These results show that brain activity in the primary visual cortex was strongly related to behavioral performance in Dutch participants, whereas brain activity in the primary auditory cortex was weakly related to behavioral performance in Japanese participants. Interestingly, we found a significant difference in the correlations of Japanese and Dutch groups in BA17L (z = 3.68, p < .01) and BA17R (z = 2.29, p < .05). We also found a significant difference in the correlations of Japanese and Dutch groups in BA41L (z = 3.67, p < .01), and significant tendency in BA41R (z = 1.80, p < .10). These results show that Japanese and Dutch participants revealed contrastive relations between behavior and brain activity in early sensory cortices.

**Fig 5.**
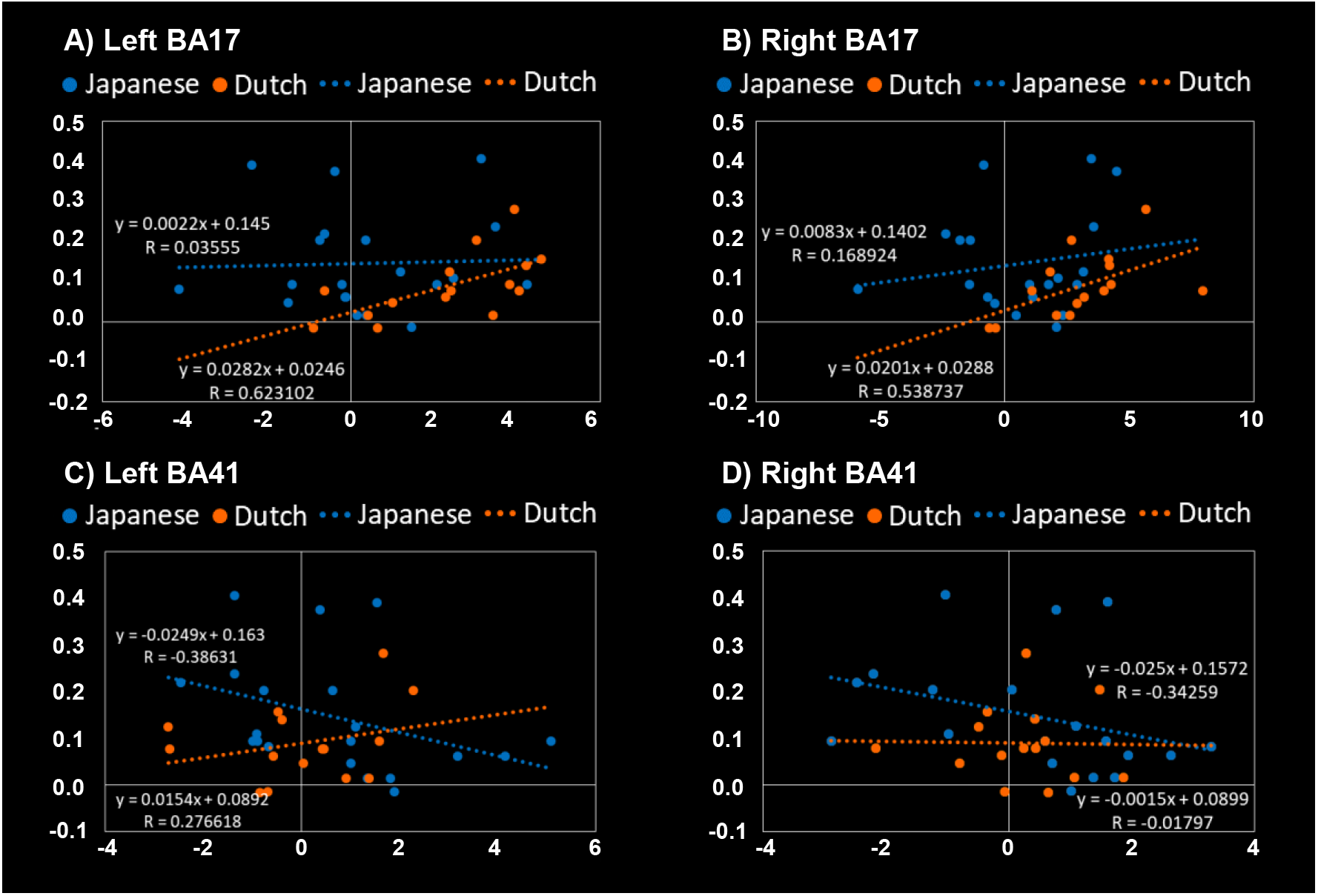
Pearson’s correlations between the congruency effects on the behavioral accuracies and those on the brain activities within the primary visual (BA17) and auditory cortices (BA41) in the face task. Effect sizes were extracted from Marsbar ROIs shown in Figure 3, and the congruency effect on the brain activities were calculated by subtracting the effect size of the incongruent condition from that of the congruent condition. In each panel, the vertical axis shows the congruency effect on behavioral accuracy while the horizontal axis shows the congruency effect on brain activity. The correlations in the left and right BA17s are shown in (A) and (B), respectively. The correlations in the left and right BA41s are shown in (C) and (D) respectively. There were significant positive correlations between the congruency effects on behavioral accuracy and those on brain activity in Dutch in both BA17L and BA17R, but not at BA41L and BA41R.

## Discussion

### Behavioral results

In this study, we investigated the neural basis of cultural differences in emotion perception from multisensory signals. Specifically, we focused on the roles of early sensory areas in the multisensory perception of emotion. In the experiment, Japanese and Dutch participants were presented with movies showing congruent or incongruent emotions expressed in faces and voices. Participants were asked to judge the emotion from the face or voice. Behavioral results showed that audiovisual consistency of stimuli affects accuracy in both face and voice tasks regardless of culture. These results are consistent with previous studies (Collignon et al., 2008; de Gelder & Vroomen, 2000; Liu et al., 2015; Takagi et al., 2015; Tanaka et al., 2010).

Behavioral results also revealed cultural differences in congruency effects. Specifically, the two-way interaction between task and group was significant, suggesting that culture affects the cross-modal bias. This interaction is consistent with Tanaka et al. (2010), which compared Japanese and Dutch participants, and Liu et al. (2015), which compared Chinese and English-speaking participants. Our results are also in line with the previously reported cultural differences in unisensory processes stating that East Asians (compared with Westerners) tend to use a different strategy in judging facial expressions (Jack et al., 2009; Yuki et al., 2007), and have a different attentional bias to different types of facial (Masuda et al., 2008) and vocal (Ishii et al., 2003) information. Our results on the less visual susceptibility in Japanese are also in line with the finding that Japanese speakers use visual information less than English speakers do in perceiving audiovisual speech (Sekiyama & Tohkura, 1991).

Note that the congruency effects were not significantly different between Japanese and Dutch groups in each task, although the task by group interaction was significant. This does not replicate the results of Tanaka et al. (2010), which found cultural differences in both face and voice tasks between Japanese and Dutch participants. As the current fMRI study was conducted in a different auditory context in the scanner, performance in the voice task was lower than those in the face task, although we set the difficulty level on the basis of the results of preliminary experiments. This difference in difficulty level might have affected the results.

### fMRI results

fMRI results showed that audiovisual consistency affects brain activity in the bilateral primary visual cortices in the face tasks, and in the bilateral primary auditory cortices in the voice tasks. These results suggest that multisensory interactions of affective faces and voices occur at a relatively low level in the cortical hierarchy. In this experiment, participants were instructed to attend to either face or voice and to ignore the other whether it is congruent or incongruent with each other. Since participants’ attention was controlled between congruent and incongruent conditions, the congruency effects in sensory cortices is considered to reflect multisensory interactions, rather than mere attentional effects. In addition to the cross-modal role of the primary sensory areas (e.g., Sadato et al., 1996), several lines of evidence suggest that primary sensory areas play an important role in multisensory interaction. Multisensory interactions in V1 are consistent with those observed in previous studies using an illusory visual flash (Watkins et al., 2006, 2007). Watkins et al. (2006) used high field fMRI to study brain activity associated with an established audiovisual illusion. When a single brief visual flash was accompanied by two auditory beeps, it was frequently perceived incorrectly as two flashes (Shams et al., 2000). They found that perception of this ‘fission’ illusion was associated with increased activity in retinotopic areas of the human primary visual cortex representing the visual stimulus. Multisensory interactions in the auditory cortex are consistent with previous studies using congruent and incongruent (McGurk type) audiovisual speech stimuli (Erickson et al., 2014; Okada et al., 2013). Okada et al. (2013) have demonstrated that congruent visual speech increases the activity in the bilateral auditory cortex. Erickson et al. (2014) have shown that McGurk stimuli increases activity in left pSTG. Effects of visual information on the activities in A1 have also been observed in studies of non-human species (Ghazanfar et al., 2005; Kayser et al., 2008; Schroeder et al., 2005; Werner-Reiss et al., 2003). Our results are consistent with these findings and extend our knowledge regarding the multisensory interactions in the visual and auditory cortices using affective audiovisual stimuli. The congruency effects in early sensory areas when judging emotions also are consistent with the idea that early sensory cortices code the valence of socio-emotional signals (Miskovic & Anderson, 2018; Shinkareva et al., 2014) and extend their argument to multisensory roles of early sensory cortices.

Importantly, our findings suggest that culture affects brain activity at early sensory levels. fMRI results revealed that the congruency effects in the bilateral primary visual cortices in the face task were larger in Dutch than in Japanese, but not in the voice task. Liu et al. (2015) compared the ERP between Chinese and English-speaking groups and showed cultural differences in N400 during semantic processing. Taken together, the impact of culture on audiovisual emotion perception occurs at both the sensory and semantic levels. Considering that the amygdala response to fearful faces is modulated by culture (Chiao et al., 2008; Harada et al., 2020), both early sensory cortices and limbic/paralimbic regions may play differential roles in emotion judgement and be affected by culture.

In contrast, the congruency effects in brain activity for both face and voice tasks were not significantly different between Dutch and Japanese in the primary auditory cortex, although the congruency effects tend to be larger in Japanese group. This pattern corresponds to the behavioral results showing that the congruency effects were not significantly different between Japanese and Dutch groups in the voice tasks. Thus, future studies should further examine this asymmetric pattern between auditory-to-visual and visual-to-auditory influences in both behavioral and neural levels.

### Correlations between brain activity and behavior

The results of correlation analyses between brain activity and behavior suggest that differential sensory areas are related to behavioral performance between Dutch and Japanese people. In Dutch participants, we found reliable positive correlations between the congruency effects in the bilateral primary visual cortices and behavioral interferences. In contrast, congruency effects in the primary auditory cortex did not correlate with behavioral interferences. Interestingly, Japanese participants revealed an opposite pattern. In Japanese participants, there was a weak negative correlation between congruency effects in the primary auditory cortex and behavioral interferences. In contrast, congruency effects in the primary visual cortex did not correlate with behavioral interferences. Importantly, the correlations between behavior and brain activities in Dutch and Japanese groups were significantly different in both visual and auditory cortices. In the face task, task-irrelevant auditory input is analyzed in the primary auditory cortex and might be conveyed to the primary visual cortex although it is not evident whether it is conveyed directly (e.g., Falchier et al., 2002), or via multisensory integration sites such as pSTS and other integration sites (Calvert et al., 2000; Wright et al., 2003; Nath and Beauchamp, 2012; Stevenson et al., 2011).

On the basis of the findings, we speculate that the results of correlation analyses can be interpreted as follows: Assuming that task-irrelevant vocal information affects the activity in V1, the Dutch results can be understood. This may be triggered by inhibitory signals sent to V1 when incongruent vocal input is detected in A1. Consequently, task-irrelevant vocal information might interfere with early visual analyses of facial expressions in V1. As they inhibit visual processing at V1 (i.e., larger congruency effects in V1), their behavioral performance in the face task would decrease (i.e., larger congruency effects at behavioral level). There are also individual differences in the efficiency of inhibition; Some Dutch participants can efficiently inhibit vocal information in V1 (i.e., smaller congruency effects in V1), which lead to better performance in the face task (i.e., smaller congruency effects at behavioral level). Other Dutch participants cannot efficiently inhibit vocal information in V1, (i.e., larger congruency effects in V1), which lead to worse performance in the face task (i.e., larger congruency effects at behavioral level).

In contrast to Dutch participants, correlations between brain activity in V1 and behavioral results were not observed in Japanese participants. This leads to the possibility that behavioral interference occurs in brain areas other than V1 in Japanese participants. In Japanese participants, task-irrelevant vocal information might be inhibited at the level of A1. There are also individual differences in the efficiency of inhibition; Some Japanese participants can efficiently inhibit vocal information in A1 (i.e., larger congruency effects in A1), which lead to better performance in the face task (i.e., smaller congruency effects at behavioral level). Other Japanese participants cannot efficiently inhibit vocal information in A1, (i.e., smaller congruency effects in A1), which lead to worse performance in the face task (i.e., larger congruency effects at behavioral level).

In sum, the results of correlation analyses suggest that V1 is related to behavioral performances in Dutch, whereas A1 is related to behavioral performances in Japanese. Thus, our results raise the possibility that differential sensory areas are related to behavioral performances in multisensory perception depending on cultures. Although these possibilities are speculative at the moment, our results yield verifiable hypotheses, which could be examined in future studies.

## Conclusions

Using fMRI, the current study is the first to report that culture affects the activities of early sensory areas in multisensory perception. fMRI results revealed that the congruency effects in the bilateral primary visual cortices in the face tasks were larger in Dutch than in Japanese. The results of correlation analyses between brain activity and behavior suggest that different sensory areas are related to behavioral performance in different cultures. These findings should guide further research on how culture shapes multisensory perception.

## Acknowledgments

This work was supported by the Cooperative Study Program (2018, No.611) of National Institute for Physiological Sciences. This work was supported by JSPS KAKENHI Grant-in-Aid for Young Scientists (A) (No. 24680030), and Grant-in-Aid for Scientific Research on Innovative Areas No. 17H06345 “Construction of the Face-Body Studies in Transcultural Conditions” (to AT). BdG and EH were supported by the European Research Council (ERC) FP7-IDEAS-ERC (Grant 15 agreement number 295673; Emobodies).

